# Genetics of Cocaine and Methamphetamine Consumption and Preference in *Drosophila melanogaster*

**DOI:** 10.1101/470906

**Authors:** Chad A. Highfill, Brandon M. Baker, Stephenie D. Stevens, Robert R. H. Anholt, Trudy F. C. Mackay

## Abstract

Illicit use of psychostimulants, such as cocaine and methamphetamine, constitutes a significant public health problem. Whereas neural mechanisms that mediate the effects of these drugs are well-characterized, genetic factors that account for individual variation in susceptibility to substance abuse and addiction remain largely unknown. *Drosophila melanogaster* can serve as a translational model for studies on substance abuse, since flies have a dopamine transporter that can bind cocaine and methamphetamine, and exposure to these compounds elicits effects similar to those observed in people, suggesting conserved evolutionary mechanisms underlying drug responses. Here, we used the *D. melanogaster* Genetic Reference Panel to investigate the genetic basis for variation in psychostimulant drug consumption, to determine whether similar or distinct genetic networks underlie variation in consumption of cocaine and methamphetamine, and to assess the extent of sexual dimorphism and effect of genetic context on variation in voluntary drug consumption. Quantification of natural genetic variation in voluntary consumption, preference, and change in consumption and preference over time for cocaine and methamphetamine uncovered significant genetic variation for all traits, including sex-, exposure-and drug-specific genetic variation. Genome wide association analyses identified both shared and drug-specific candidate genes, which could be integrated in genetic interaction networks. We assessed the effects of ubiquitous RNA interference (RNAi) on consumption behaviors for 34 candidate genes: all affected at least one behavior. Finally, we utilized RNAi knockdown in the nervous system to implicate dopaminergic neurons and the mushroom bodies as part of the neural circuitry underlying experience-dependent development of drug preference.

**AUTHOR SUMMARY:** Illicit use of cocaine and methamphetamine is a major public health problem. Whereas the neurological effects of these drugs are well characterized, it remains challenging to determine genetic risk factors for substance abuse in human populations. The fruit fly, *Drosophila melanogaster*, presents an excellent model for identifying evolutionarily conserved genes that affect drug consumption, since genetic background and exposure can be controlled precisely. We took advantage of natural variation in a panel of inbred wild derived fly lines with complete genome sequences to assess the extent of genetic variation among these lines for voluntary consumption of cocaine and methamphetamine and to explore whether some genetic backgrounds might show experience-dependent development of drug preference. The drug consumption traits were highly variable among the lines with strong sex-, drug- and exposure time-specific components. We identified candidate genes and gene networks associated with variation in consumption of cocaine and methamphetamine and development of drug preference. Using tissue-specific suppression of gene expression, we were able to functionally implicate candidate genes that affected at least one consumption trait in at least one drug and sex. In humans, the mesolimbic dopaminergic projection plays a role in drug addiction. We asked whether in Drosophila the mushroom bodies could play an analogous role, as they are integrative brain centers associated with experience-dependent learning. Indeed, our results suggest that variation in consumption and development of preference for both cocaine and methamphetamine is mediated, at least in part, through a neural network that comprises dopaminergic projections to the mushroom bodies.

## INTRODUCTION

Illicit use of cocaine and methamphetamine constitutes a significant public health problem that incurs great socioeconomic costs in the United States and worldwide [1-3]. Cocaine and the amphetamine class of drugs are potent central nervous system stimulants that act by raising synaptic concentrations of biogenic amines. Cocaine inhibits neurotransmitter reuptake at dopaminergic, serotonergic and noradrenergic synapses [4,5]. Amphetamine increases neurotransmission by promoting the release of dopamine from presynaptic vesicles through its actions on the vesicular monoamine transporter and subsequent reverse flux of dopamine via the dopamine transporter and through the plasma membrane into the synaptic cleft [6,7].

Amphetamine, methamphetamine, and methylphenidate are used clinically to treat attention deficit hyperactivity disorder and narcolepsy. Long term use of these compounds, however, can lead to addiction, and ultimately death [8]. The addictive properties of these drugs are mediated through the dopaminergic mesolimbic reward pathway, which projects from the ventral tegmental area via the nucleus accumbens to prefrontal cortex [9]. Although most studies on psychostimulants focus on addiction, addiction represents only one facet of the diverse organismal effects that result from psychostimulant drug abuse. These drugs exert a wide range of physiological and behavioral effects, including suppression of appetite, which can result in malnutrition, and severe cardiovascular, respiratory and renal disorders. Use of cocaine and amphetamine can also cause mental disorders, including paranoia, anxiety, and psychosis [10,11].

Susceptibility to the effects of cocaine and methamphetamine is likely to vary among individuals and be determined both by environmental and genetic factors. However, there is limited information regarding the genetic basis of susceptibility to the effects of these drugs in human populations [12]. Twin and adoption studies have focused primarily on alcohol abuse and illicit drugs, such as cannabis, with heritability estimates ranging from ~30-70% [13,14]. Most studies on psychostimulant addiction to date have centered on candidate genes associated with neurotransmission in the mesolimbic projection [12], and many of these are inconclusive or contradictory. For example, some studies reported that alleles of the dopamine D2 receptor were associated with substance abuse [15-18], whereas others did not replicate this finding [19-24]. Similar contradictory results have been obtained for association analyses between polymorphisms in the dopamine transporter gene and cocaine-related phenotypes [24-28]. These contradictory findings may be due in part to failure to account for multiple testing or population structure [29]. However, genetic studies of substance abuse and addiction in human populations are challenging due to diverse social conditions and physical environments, confounding factors with comorbid conditions such as alcoholism or psychiatric disorders, and difficulty to recruit large numbers of study subjects due to criminalization.

*Drosophila melanogaster* is an excellent model for identifying genes that affect drug consumption behaviors since both the genetic background and environment, including exposure to drugs, can be controlled precisely. These results have translational potential since 75% of disease-causing genes in humans have a fly ortholog [30]. High resolution X-ray crystallography has shown that the *D. melanogaster* dopamine transporter has a central conformationally pliable binding site that can accommodate cocaine, methamphetamine and their closely related analogues [31]. Similar to its effects in humans, methamphetamine suppresses sleep, causes arousal and suppresses food intake in flies [32-34]. In addition, cocaine, amphetamine and methylphenidate exert quantifiable locomotor effects in flies [35-41]. Thus, despite profound differences between the neuroanatomical organization of the fly and vertebrate brains, it is likely that behavioral and physiological effects of methamphetamine and cocaine are mediated, at least in part, by evolutionarily analogous mechanisms.

Here, we used the inbred, sequenced lines of the *D. melanogaster* Genetic Reference Panel (DGRP [42,43]) to investigate the genetic basis for variation in psychostimulant drug consumption. We used a two-capillary Capillary Feeding (CAFE) assay [44-46] to quantify voluntary consumption, preference and change of consumption and preference over time for cocaine and methamphetamine. Since cocaine and methamphetamine both target dopaminergic synaptic transmission, but through different mechanisms, we asked to what extent genetic networks that underlie variation in consumption of cocaine and methamphetamine incorporate the same or different genes. We also sought to determine the extent of sexual dimorphism for naïve and experience-dependent voluntary drug intake. In addition, we asked how much variation in voluntary drug consumption exists among different DGRP lines and what fraction of that variation is accounted for by genetic variation. We showed that there is naturally occurring genetic variation for all drug consumption traits with strong sex-, drug- and exposure time-specific components. We performed genome wide association (GWA) analyses to identify candidate genes associated with the drug consumption behaviors that could be mapped to a genetic interaction network. We tested the effects of RNAi mediated suppression of gene expression [47] on all consumption behaviors for 34 candidate genes and found that all affected at least one behavior in at least one drug and sex. Finally, we used RNAi to suppress gene expression in neurons, glia, the mushroom bodies and dopaminergic neurons in a subset of genes and showed that innate preference and the development of preference for psychostimulant drugs involves dopaminergic neurons and the mushroom bodies, neural elements associated with experience-dependent modulation of behavior.

## RESULTS

### Quantitative genetic analysis of drug consumption behaviors in the DGRP

We used a two-capillary CAFE assay [44-46] to enable flies to choose to consume either sucrose or sucrose supplemented with 0.2 mg/ml cocaine (or 0.5 mg/ml methamphetamine), analogous to the two-bottle choice assay used in rodent studies [48] (Fig.1). We quantified consumption for three consecutive days for males and females from each of 46 DGRP lines that were unrelated, free of chromosomal inversions, and free of infection with the endosymbiont *Wolbachia pipientis* [43; Table S1]. These data enabled us to assess whether there is naturally occurring genetic variation in this population for naïve consumption of each solution and preference, and change of consumption and preference upon repeated exposures (*i.e*., experience-dependent modification of behavior).

**Fig. 1.**
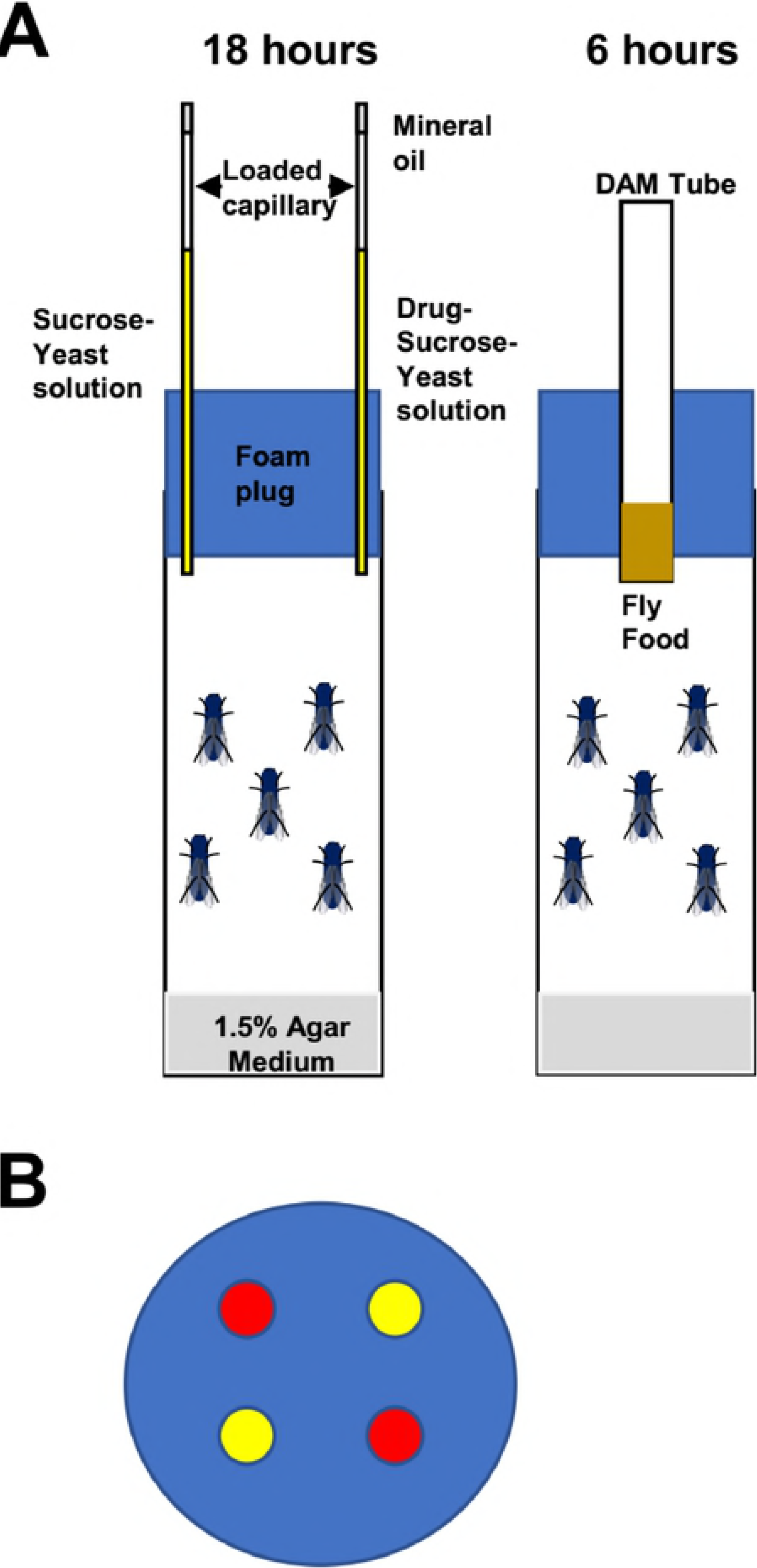
Consumption and preference assay. (A) Cartoon illustrating the four capillary CAFÉ assay. Each of the three exposures consists of an 18 hour feeding trial with sucrose or drug + sucrose, followed by 6 hours recovery with standard culture medium. (B) Positions of capillaries with the two solutions (indicated by red and yellow).

We performed four-way mixed model analyses of variance (ANOVA) to partition variation in consumption between DGRP lines, males and females, drug *vs*. sucrose, and the three exposures. All main effects were significant for both drugs (Table 1), indicating genetic variation for consumption, difference between amount of sucrose and drug consumed, sexual dimorphism, and experience-dependent modulation of behavior. We are most interested in the two- and three-way interaction terms involving Line, as they indicate genetic variation in sexual dimorphism (*L*×*X*), change of consumption between exposures (*L*×*E*), preference for sucrose or drug solution (*L*×*S*), and change of preference for sucrose or drug between exposures (*L*×*E*×*S*). With the exception of *L*×*S*, these interaction terms were significant for both the cocaine and methamphetamine analyses (Table 1).

**Table 1.**
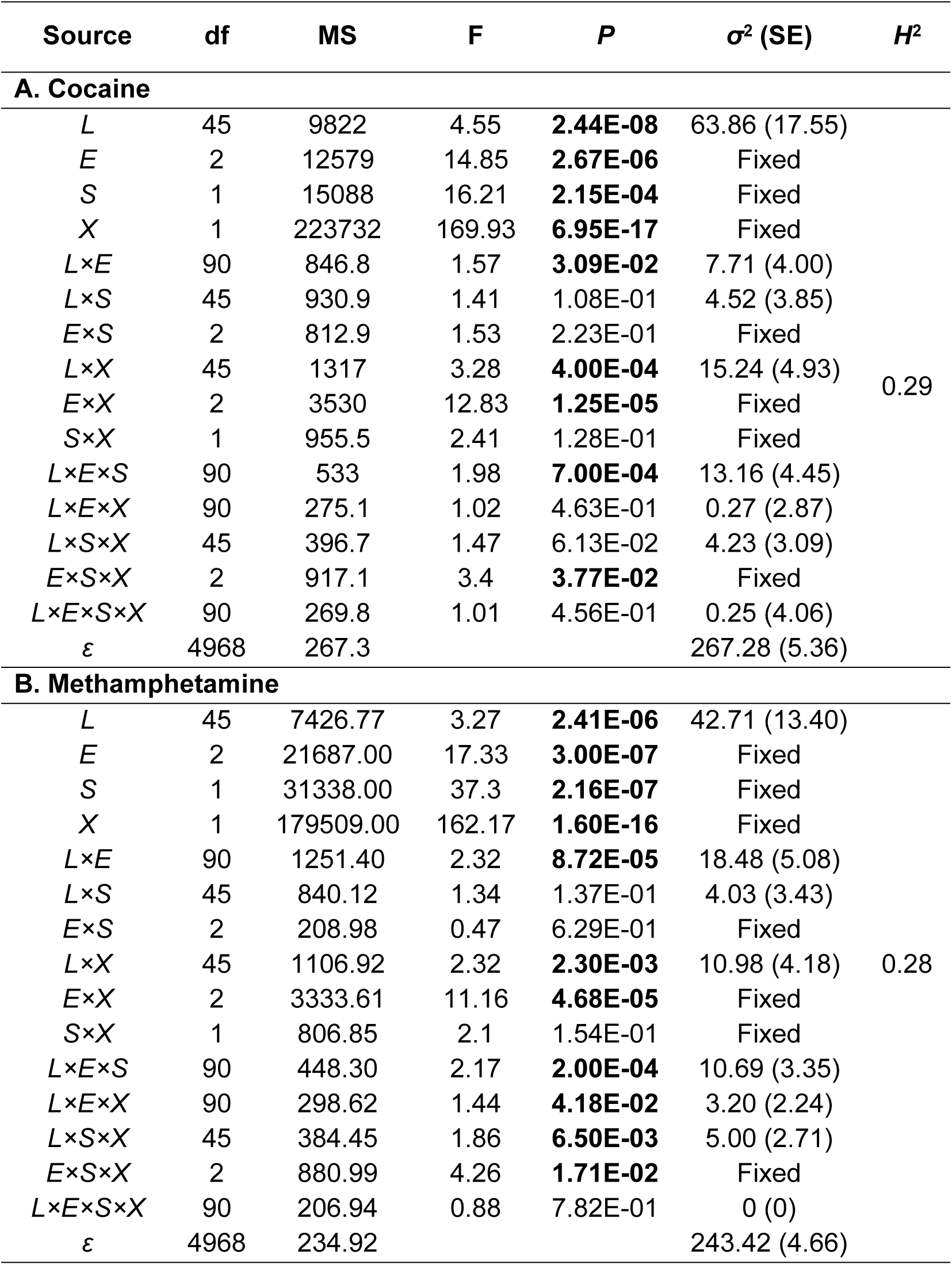
Analyses of variance of consumption measured over three exposures. Exposure, Sex, Solution, and their interaction are fixed effects, the rest are random. E: Exposure; X: Sex; S: Solution; L: DGRP Line; ε: residual; df: degrees of freedom; MS: Type III mean squares; F: F-ratio test; P: P-value; σ2: variance component estimate; SE: standard error; H2: Broad sense heritability. Significant P-values are highlighted in bold font.

We next performed reduced ANOVA models to quantify broad sense heritabilities (*H*^2^) for consumption and change in consumption traits (Table S2). We found significant genetic variation in consumption of both drugs and sucrose alone within each sex and exposure, with *H*^2^ ranging between 0.20 and 0.38 for cocaine consumption and between 0.22 and 0.30 for methamphetamine consumption (Fig. 2, Table S2). Further, there was significant genetic variation for the change in consumption of sucrose alone or drug in both sexes between the third and first exposures, with *H*^2^ ranging between 0.14 and 0.18 for cocaine and between 0.17 and 0.22 for methamphetamine (Fig. 2, Table S2). Thus, there is genetic variation for both consumption and experience-dependent consumption of both drugs and sucrose alone in the DGRP.

**Fig. 2.**
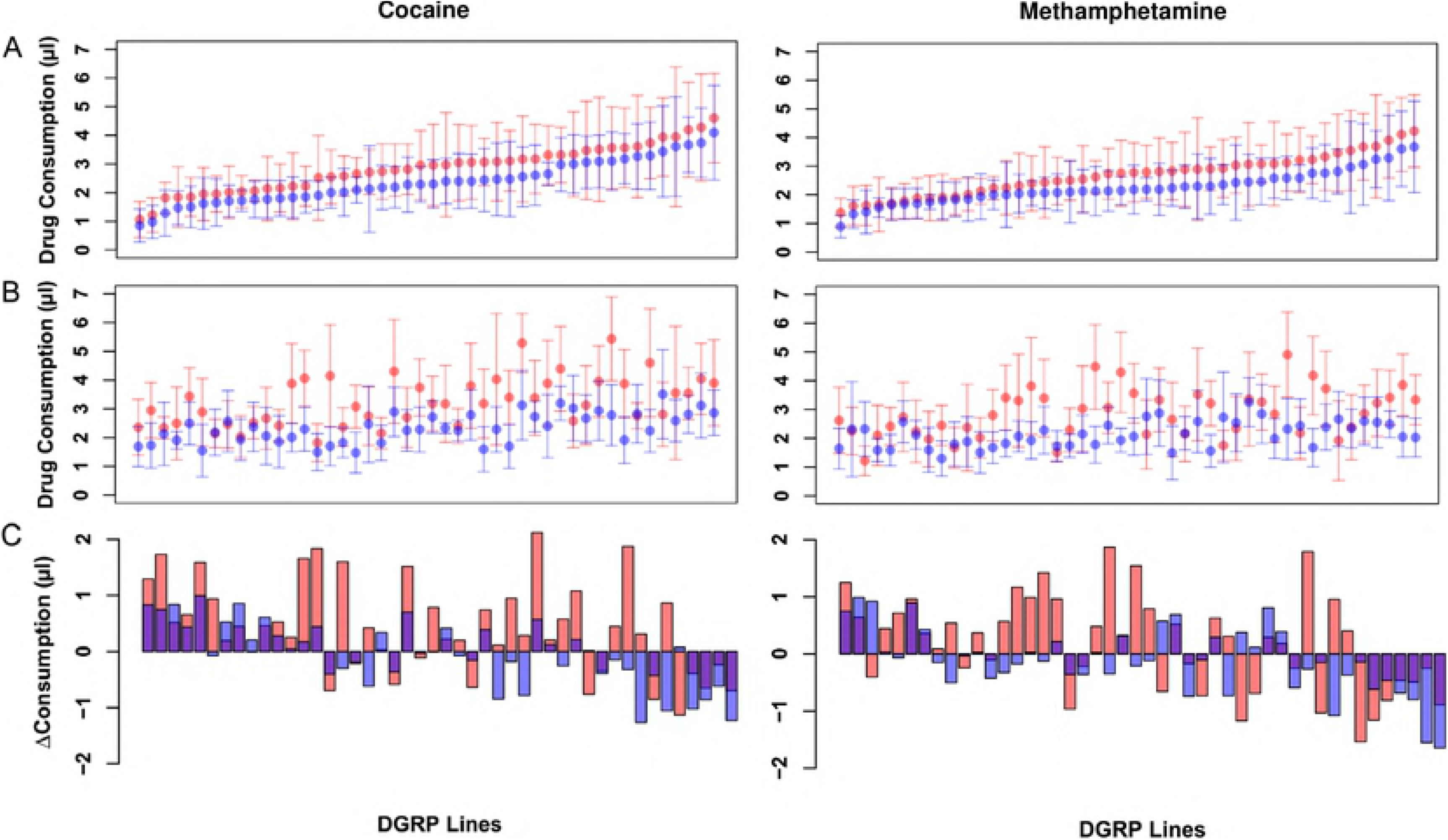
Variation in drug consumption among 46 DGRP lines. (A) Initial exposure. Lines are from lowest to highest consumption in females. (B) Third exposure. The line order is the same as in (A). (C) Change in consumption between exposures 3 and 1. Positive values indicate increased drug consumption in Exposure 3. The line order is the same as in (A). Pink denotes females, blue indicates males, and purple is overlap of both sexes. Error bars are ± 1SD.

Finally, we defined preference in two ways: as the difference between amount of drug and sucrose alone consumed (Preference A), and as this difference scaled by the total amount of both solutions consumed (Preference B). Preference values of 0 indicate equal consumption of sucrose alone and sucrose containing drug; values > 0 represent preference for the drug and values < 0 indicate drug avoidance. Both preference metrics were significantly genetically variable for each sex and exposure for cocaine, with *H*^2^ ranging from 0.06-0.16; while for methamphetamine, both preference metrics were significantly genetically variable in females for all exposures (*H*^2^ from 0.05-0.18) and for males in the second and third exposures (*H*^2^ from 0.08-0.11) (Table S2). For cocaine, the difference in preference A between exposures 3 and 1 was significant only in females (*H*^2^ = 0.11) while the difference in Preference B was significant for females (*H*^2^ = 0.13) and males (*H*^2^ = 0.05). For methamphetamine, the difference in Preference A was significant in males (*H*^2^ = 0.04) and the difference in Preference B was significant in females (*H*^2^ = 0.04) (Table S2). Thus, there is genetic variation for both innate drug preference and experience-dependent drug preference in the DGRP.

The heritabilities of consumption traits are low, as is typical for behavioral traits, indicating that environmental factors, including previous experience, predominantly contribute to the observed phenotypic variation. The advantage of performing multiple replicate measurements of each DGRP line is that the broad sense heritabilities of line means (Table S3) used in the GWA analyses (see below) are much greater than heritabilities based on individual vial replicates (Table S2).

We computed the genetic and phenotypic correlations between males and females for the consumption behaviors, between exposures for consumption and preference, and between solutions (Table S4). Cross-sex genetic correlations for consumption tended to decrease with the number of exposures for both cocaine and methamphetamine, suggesting that the experience-dependent modification of consumption is sex-specific. Consumption of drugs and sucrose is highly correlated across the three exposures (albeit significantly different from unity), while the correlations of drug preference across exposures are low to moderate for both cocaine and methamphetamine in both sexes. Although the consumption of drugs and sucrose for cocaine and methamphetamine are genetically and phenotypically correlated in both sexes, preference for the two drugs is not significantly correlated. Finally, Preference A and Preference B within each exposure are nearly perfectly correlated, as expected since the difference in consumption is in both metrics.

In summary, we found that there is extensive genetic variation in consumption and preference as well as change in consumption and preference with repeated exposures for both cocaine and methamphetamine across different genetic backgrounds, and that genetic variation for these traits has significant sex- and drug-specific components.

### Genome wide association analyses of drug consumption in the DGRP

Our quantitative genetic analyses of consumption in the DGRP indicate that there is genetic variation for all traits assessed, and that the traits have a complex correlation structure indicating partially common and partially distinct genetic bases. Therefore, we performed single variant GWA analyses for 12 traits (drug and sucrose consumption exposure 1, drug and sucrose consumption exposure 3, change in drug and sucrose consumption, preference A exposure 1, preference A exposure 3, preference B exposure 1, preference B exposure 3, change in preference A, and change in preference B) for cocaine and methamphetamine, separately for males and females. We performed association tests for 1,891,456 DNA sequence variants present in the 46 DGRP lines with minor allele frequencies greater than 0.05 [43].

At a lenient significance threshold of *P* < 5 × 10^-5^, we identified 1,441 polymorphisms in or near (within 1 kb of the start and end of the gene body) 725 genes for all consumption behaviors related to cocaine, and 1,413 polymorphisms in or near 774 genes for methamphetamine exposure (Table S5). The majority of these variants had sex-specific effects. A total of 40 variants and 141 genes overlapped between cocaine and methamphetamine. The variants in or near genes implicate candidate genes affecting consumption behaviors, while the intergenic variants could potentially contain regulatory motifs for transcription factor-binding sites or chromatin structure regulating these traits. Only two variants are formally significant following a Bonferroni correction for multiple tests (*P* < 2.64 × 10^−8^). *2L*_10179155_SNP is located within an intronic region in *CG44153* and affects experience-dependent development of methamphetamine preference in both sexes. Its human homolog *ADGRB3* encodes a G-protein coupled receptor, which contributes to the formation and maintenance of excitatory synapses [49] and has been implicated in GWA studies on human addiction [50]. *3R*_27215016_SNP is a synonymous SNP in the coding sequence of *CG1607* and affects naïve consumption of sucrose. *CG1607* encodes an amino acid transmembrane transporter. One of its human orthologs, *SLC7A5*, is an amino acid transporter, mutations in which are associated with autism spectrum disorder and defects in motor coordination [51].

While not formally significant, we identified genes previously associated with cocaine-related behaviors (*Bx* [*Lmo*], *loco*, *Tao*) and ethanol-related behaviors (*Bx*, *DopR*, *Egfr*, *hppy*, *Tao*, *Tbh*) [52] in *D. melanogaster*. In addition, the genes implicated by the GWA analyses are enriched for multiple gene ontology (GO) categories and pathways [53,54] at a false discovery rate < 0.05 (Table S5). GO terms involved in nervous system development and function were among the most highly enriched, consistent with the known neurobiological mechanisms of action of these drugs. Finally, we note that ~ 70% of the candidate genes from the GWA analyses have human orthologs, and many of these genes have previously been associated with cocaine or methamphetamine abuse in humans or with behaviors associated with intake and response to various psychoactive substances (alcohol, cannabis, nicotine, opioids) in humans as well as zebrafish, mouse and rat models (Table S6). This suggests that cocaine and methamphetamine exert their effects in flies and humans through evolutionarily conserved neural mechanisms.

These results suggest a highly polygenic architecture for variation in consumption and drug preference, and that the genetic underpinnings for variation in consumption or preference are both shared and distinct for cocaine and methamphetamine, consistent with the quantitative genetic analyses.

### A genetic interaction network for consumption behaviors

We next asked whether the genes we identified in the GWA analyses belonged to a known genetic interaction network. Since the consumption behaviors are highly inter-correlated, we queried whether all 1,358 candidate genes from the GWA analyses for both cocaine and methamphetamine combined could be clustered into significant sub-networks based on curated genetic interactions in Drosophila. If we do not allow any missing genes, we find a significant (*P* = 9.99 × 10^-4^) network of 81 candidate genes (Fig. 3, Table S7), most of which (88.9%) are predicted to have human orthologs [55].

**Fig. 3.**
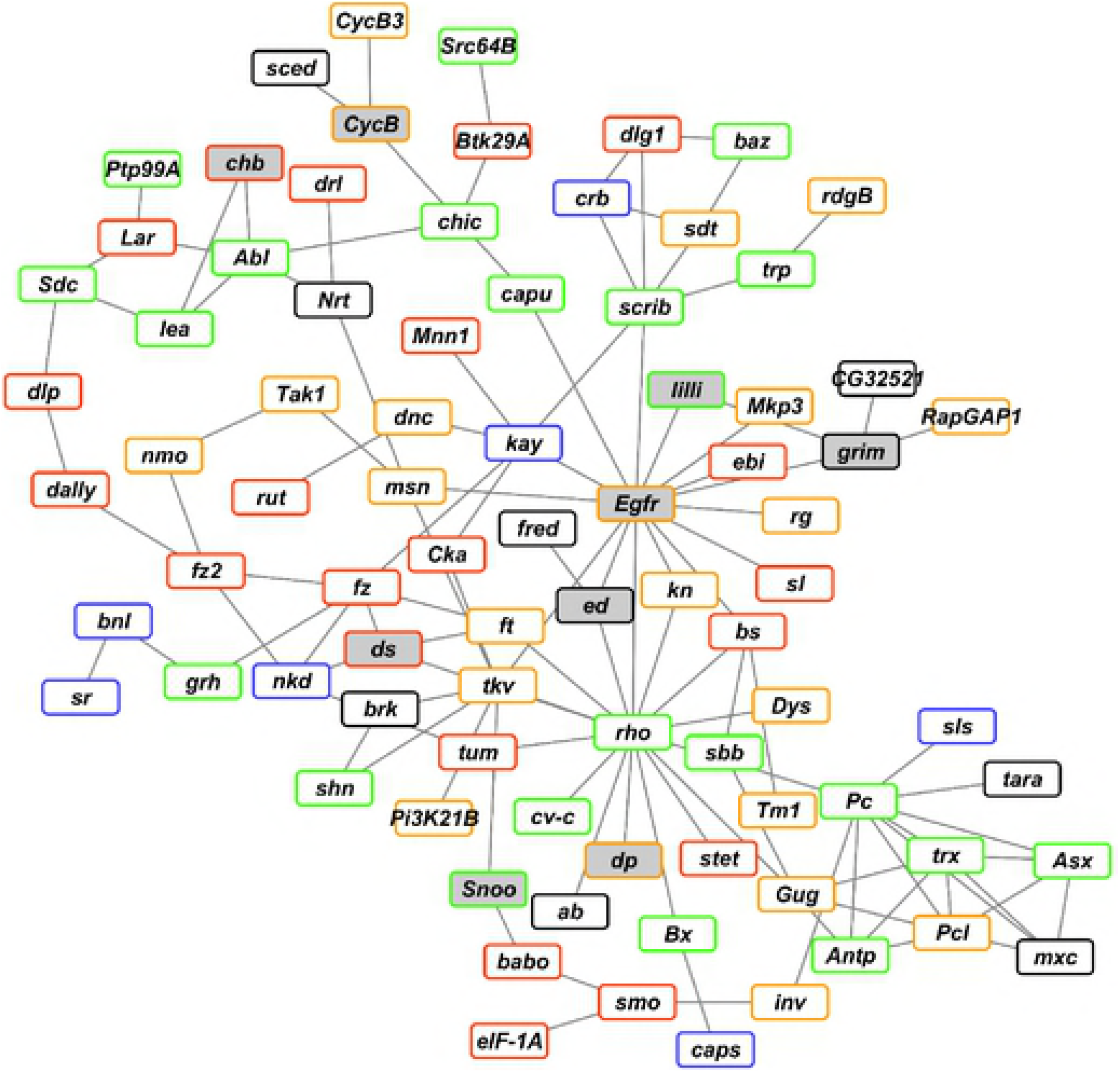
Significant genetic interaction network of genes identified in the GWA analyses for all cocaine and methamphetamine related traits combined. Borders indicate the strength of the evidence for a human ortholog. Black: DIOPT score < 3; Blue: DIOPT score 3-6; Green: DIOPT score 7-9; Orange: DIOPT score 10-12; Red: DIOPT score 13-15. Grey boxes have effects on at least one drug-seeking behavior from RNAi knockdown of gene expression.

**Fig. 4.**
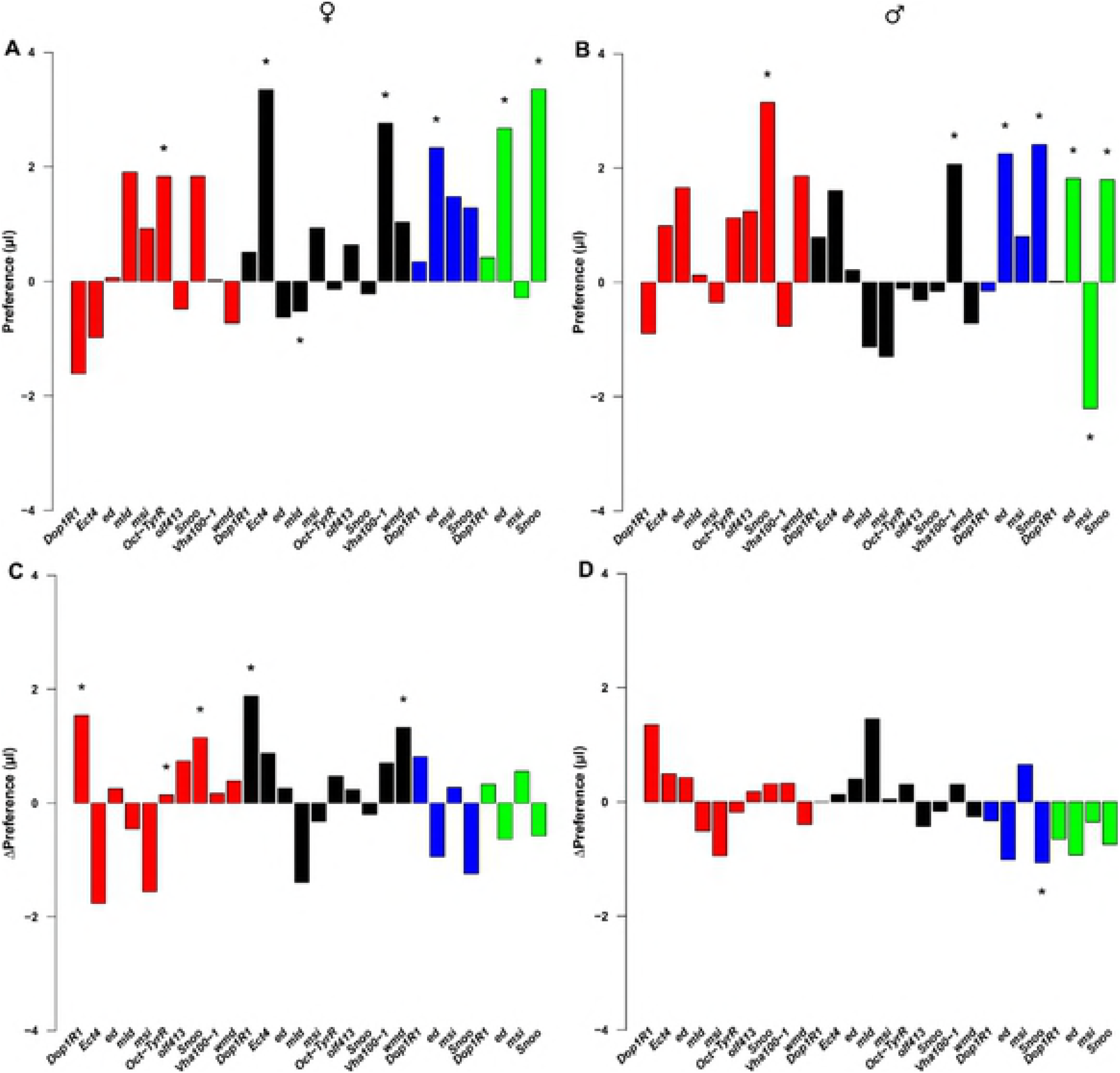
Differences in cocaine preference and change in cocaine preference between the third and first exposures between RNAi and control genotypes. (A) Female preference. (B) Male preference. (C) Female change of preference. (D) Male change of preference. Red, black, blue, and green bars denote *elav-GAL4, repo-GAL4, 201Y-GAL4* and *TH-GAL4* drivers, respectively. Asterisks represent significant *L*×*S* terms (A, B) or significant *L*×*S*×*E* terms from the full ANOVA models. Exact *P*-values are given in Table S11.

**Fig. 5.**
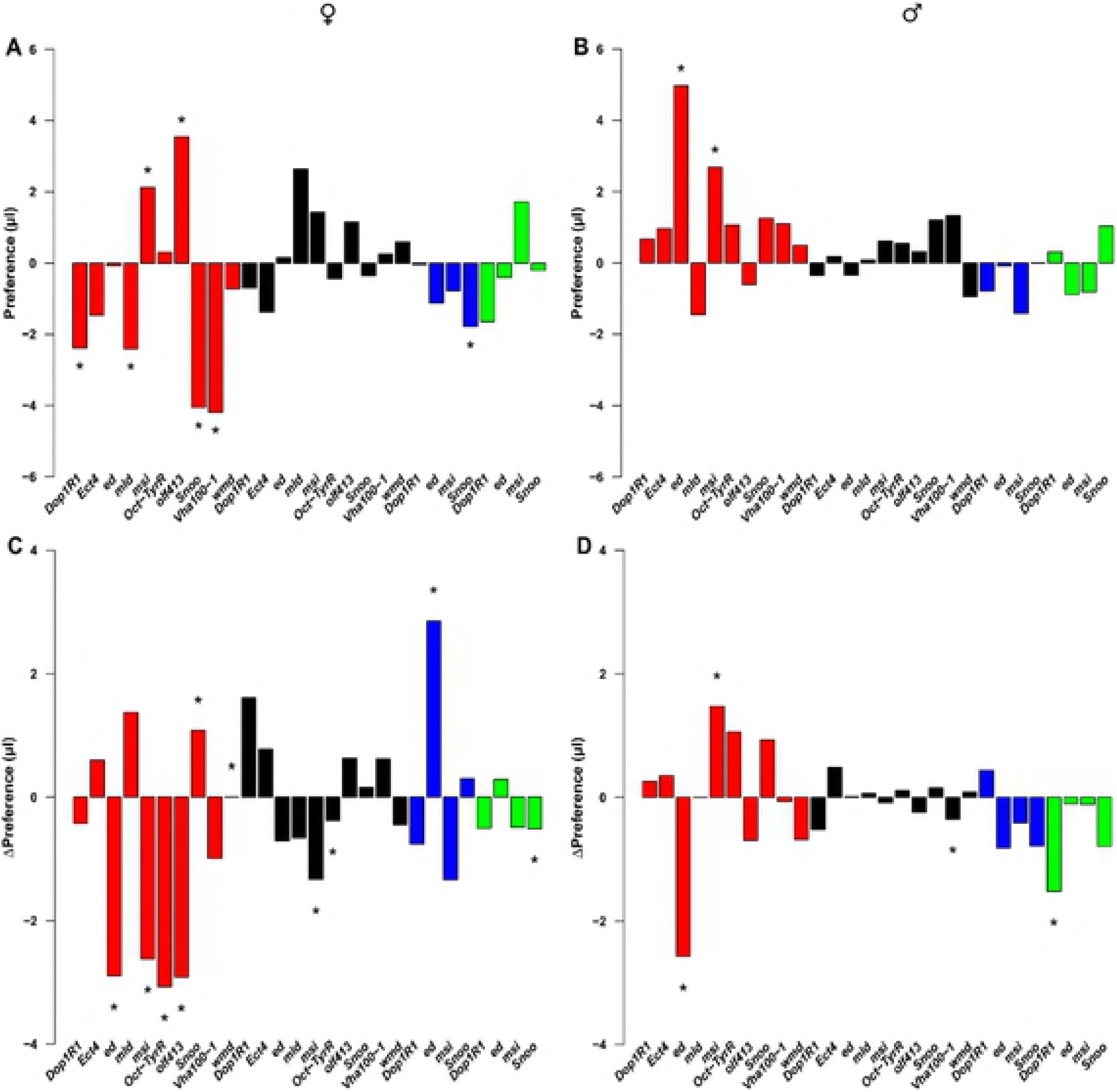
Differences in methamphetamine preference and change in methamphetamine preference between the third and first exposures between RNAi and control genotypes. (A) Female preference. (B Male preference. (C) Female change of preference. (D) Male change of preference. Red, black, blue, and green bars denote *elav-GAL4, repo-GAL4, 201Y-GAL4* and *TH-GAL4* drivers, respectively. Asterisks represent significant *L*×*S* terms (A, B) or significant *L*×*S*×*E* terms from the full ANOVA models. Exact *P*-values are given in Table S12.

We performed enrichment analyses [53,54] to gain insight in the biological context for genes in the network using a false discovery rate < 0.05. Surprisingly, many canonical signaling pathways are highly enriched, including the Wingless (Wnt), Cadherin, Cholecystokinin Receptor (CCKR), Transforming Growth factor beta (TGF), and Fibroblast Growth Factor (FGF) signaling pathways. Concomitantly, we find high enrichment of molecular function GO terms associated with regulation of transcription and DNA and protein binding, and biological function GO terms associated with development (including the development of the nervous system; Table S7). These results suggest that naturally occurring genetic variation in nervous system development is associated with variation in propensity to consume psychostimulant drugs. Furthermore, our results indicate that natural variants in key genes regulating all aspects of fly development and function can be associated with variation in drug consumption behaviors.

### Functional evaluation of candidate genes

We used RNA interference (RNAi) to functionally test whether reduced expression of candidate genes implicated by the GWA analyses affect consumption phenotypes. We selected 34 candidate genes for RNAi mediated suppression of gene expression. A total of nine of the candidate genes were in the network; the others were chosen based on gene expression in the nervous system and their known role in nervous system function, as well as belonging to enriched pathways and gene ontology categories. We measured consumption of cocaine and sucrose (Table S8) and methamphetamine and sucrose (Table S9) for three consecutive days, separately for males and females, for each of the RNAi and control genotypes, exactly as described for the DGRP lines.

We performed three-way fixed effect ANOVAs for each *UAS*-RNAi and control genotype, separately for males and females (Tables S10, S11). The main effects in these models are genotype (*L*, RNAi and control), solution (*S*, sucrose and drug) and exposure (*E*, first and third). A significant L effect denotes a difference in overall consumption between the RNAi and control genotypes; a significant *S* effect indicates a difference in preference between sucrose alone and sucrose with drug; and a significant E effect indicates a difference in consumption between exposures 1 and 3. Significant *L*×*S* and *L*×*E* interaction terms denote, respectively, a difference in preference between the RNAi and control genotypes, and a difference in consumption between exposures 1 and 3 between the two genotypes. A significant *L*×*S*×*E* interaction indicates a change in preference with repeated exposure between the RNAi and control genotypes. We are most interested in the main effect of genotype and interactions with genotype; *i.e.*, consumption, preference, change of consumption and change of preference.

First, we used a weak ubiquitous *GAL4* driver crossed to all 34 *UAS*-RNAi genotypes and their respective controls. All candidate genes had a significant (*P* < 0.05) effect on at least one of the consumption traits in at least one drug or sex combination. A total of 22 (25) genes affected consumption of cocaine (methamphetamine), 21 (23) affected a change of consumption with exposure to cocaine (methamphetamine), 16 (10) affected cocaine (methamphetamine) preference, and 11 (11) affected a change in cocaine (methamphetamine) preference with exposure in males and/or females (Tables S10, S11, Fig.s S1-S3). There were pronounced sex- and drug-specific effects for all drug-related traits. The majority of RNAi genotypes showed reduced consumption of cocaine and/or methamphetamine compared to their controls, dependent on exposure and sex. If consumption is positively associated with gene expression, this suggests that the products of these genes contribute to drug consumption. On the other hand, several RNAi constructs caused increased drug consumption, suggesting that naturally occurring variants that decrease expression of these genes could predispose to drug preference. Finally, several RNAi-targeted genes exhibit a relative increase or decrease in drug consumption compared to the control at the third exposure, indicating experience-dependent change in preference.

To extend and refine our RNAi analysis, we next selected 10 genes (*Dop1R1*, *Ect4*, *ed*, *mld*, *msi*, *Oct-TyrR*, *olf413*, *Snoo*, *Vha100-1*, *wmd*) from among those that showed phenotypic effects when targeted by RNAi under the ubiquitous driver and which have known effects on the nervous system. We assessed functional effects of these genes on consumption traits when their corresponding RNAi constructs were expressed under the control of the neuronal-specific *elav* driver or glial-specific *repo* driver. All of these genes had a significant (*P* < 0.05) effect on at least one of the consumption traits in at least one drug or sex combination under the *elav* driver, and all but *Snoo* had significant effects on at least one of the consumption traits in at least one drug or sex combination under the *repo* driver. With neuronal-specific suppression of gene expression, 9 (10) genes affected consumption of cocaine (methamphetamine), 6 (7) affected a change in consumption with exposure to cocaine (methamphetamine), 2 (7) affected cocaine (methamphetamine) preference, and 3 (6) affected a change in cocaine (methamphetamine) preference with exposure in males and/or females (Tables S10, S11, Fig. s4, S4). With glia-specific suppression of gene expression, 4 (7) genes affected consumption of cocaine (methamphetamine), 7 (6) affected a change in consumption with exposure to cocaine (methamphetamine), 3 (0) affected cocaine (methamphetamine) preference, and 2 (3) affected a change in cocaine (methamphetamine) preference with exposure in males and/or females (Tables S10, S11, Fig.s 4, 5, S4). These effects were largely sex-, drug- and driver-specific. We infer from these results that variation in gene expression in both neurons and glia contributes to phenotypic variation in drug intake behaviors.

In humans, the mesolimbic dopaminergic projection plays a role in drug addiction. In Drosophila, the mushroom bodies could play an analogous role, as they are integrative centers in the fly brain associated with experience-dependent learning [56,57], dependent on dopaminergic input. To test whether the mushroom bodies and dopaminergic projection neurons could serve as neural substrates that contribute to variation in drug consumption or preference, we focused on four genes (*Dop1R1*, *ed*, *msi*, *Snoo*,) that showed robust phenotypic effects when targeted with a corresponding *elav-*driven RNAi. Knockdown of all four genes with a mushroom body specific driver resulted in significant effects on consumption of cocaine and/or methamphetamine for at least one drug and sex combination (Tables S10, S11, Fig.s 4, 5, S5). Expression of RNAi in mushroom bodies affected change in consumption of cocaine and methamphetamine for *Dop1R1*; cocaine preference and change of methamphetamine preference for *ed*; change in consumption of cocaine for *msi*; and cocaine and methamphetamine preference, cocaine preference, change of cocaine preference and change of consumption of methamphetamine for *Snoo*. Expression of RNAi in dopaminergic neurons affected change of consumption of cocaine and change in methamphetamine preference for *Dop1R1*; consumption for cocaine and methamphetamine, change of consumption of methamphetamine and cocaine preference for *ed*; consumption of cocaine and methamphetamine, change of consumption of cocaine, and cocaine preference for msi; and all four traits for *Snoo* (Tables S10, S11, Fig.s 4, 5, S5). These effects are largely sex-, drug- and driver-specific.

These results suggest that, despite differences in the genetic underpinnings of susceptibility to cocaine and methamphetamine, phenotypic manifestation of genetic variation in consumption and development of preference for both drugs is channeled through a neural network that comprises dopaminergic projections to the mushroom bodies.

## DISCUSSION

Although studies using mice [58,59], rats [60,61], primates [62] and humans [63] provide important information about the cellular, developmental, physiological, and behavioral effects of psychostimulants, these systems are less suited to dissecting the relationship between naturally occurring genetic variation and phenotypic variation in individual susceptibility to drug consumption and/or preference. Here, we show that *D. melanogaster* harbors substantial naturally occurring variation for all consumption-related behaviors, including experience-dependent change in consumption, innate drug preference and experience-dependent change in preference, under conditions where we can obtain replicated measurements of consumption for each genotype in a choice assay performed over three successive days under controlled environmental conditions. We show that genetic variation for consumption and preference metrics is both shared between males and females and the different exposures, but is also sex-, exposure- and drug-specific. Sex differences in drug self-administration and addiction have also been shown in humans and mammalian animal models [64-72].

The Diagnostic and Statistical Manual of Mental Disorders, Fifth Edition (DSM-V) defines 11 criteria for substance use disorder in humans, all related to continuing to use of the substance despite adverse social and physiological effects and the development of tolerance with repeated exposure. The DSM-V also recognizes that there is individual variability of unknown etiology for the propensity both to experiment with psychostimulants and to develop symptoms of substance abuse following initial exposure. Previous studies of effects of cocaine [35,37-39,73-76] and methamphetamine [77] in Drosophila examined mutations and pharmacological interventions using locomotor-based assays, clearly demonstrating an adverse effect of these substances. However, previous Drosophila studies have not assessed naturally occurring variation in drug self-administration and change in this behavior on repeated exposure, which may better model the genetic basis of individual susceptibility – or resistance – to substance abuse and the development of tolerance (increased drug preference over time).

To begin to understand the nature of the genetic basis for variation in drug consumption and preference, we performed GWA analyses for all consumption traits, separately for cocaine and methamphetamine, using 1,891,456 DNA sequence variants present in the 46 DGRP lines with minor allele frequencies greater than 0.05 [43]. We identified 1,358 unique candidate genes using a lenient significance threshold of 5 × 10^-5^. We hypothesized that these candidate genes would be enriched for true positive associations despite the low power of the GWA analyses and that choosing genes for functional evaluation from this list would be more productive than choosing genes at random. Observations supporting this hypothesis are that mutations in several candidate genes have previously been shown to affect cocaine or ethanol-related phenotypes in Drosophila [52], that the candidate genes are highly enriched for GO terms involved in the development and function of the nervous system, and that 81 candidate genes can be assembled into a known genetic interaction network (Fig. 3), which is highly unlikely (*P* = 9.9 × 10^-3^) to occur by chance. The candidate genes in the significant genetic interaction network are enriched for several canonical signaling pathways as well as all aspects of development, including nervous system development. These observations suggest that subtle genetic variation in nervous system development is associated with variation in propensity for consumption of psychostimulant drugs. Nearly 90% of the genes in the network have human orthologs and are candidates for future translational studies.

We selected nine candidate genes in the significant genetic network and 25 additional candidate genes to assess whether RNAi reduction using a weak ubiquitous *GAL4* driver affected consumption traits, using the same experimental design as for the DGRP lines. All of these genes affected at least one consumption trait/sex/drug. However, there is considerable variation in the effects of different drivers on consumption, preference and change in preference for cocaine and methamphetamine, which likely reflects variation in the effects of RNA interference on different neural elements of a complex integrated neural circuitry. Indeed, several candidate genes, functionally implicated by RNAi, are associated with neural development and represent several early developmental signaling pathways. *Snoo* has been identified as a negative regulator of the decapentaplegic signaling pathway [78,79] and has been implicated in dendritic patterning [80]. Echinoid, the gene product of *ed*, is an immunoglobulin domain containing membrane protein of adherens junctions that interacts with multiple developmental signaling pathways, including Egfr, Notch and Hippo signaling [81-83]. Musashi, encoded by *msi*, is a neural RNA binding protein that interacts with Notch signaling to determine cell fate [84]. RNAi targeting of expression of these genes under *MB-GAL4* or *TH-GAL4* drivers show different effects on consumption, change in consumption, preference and change in preference for the two drugs (Fig. S5).

Among the functionally validated candidate genes, *Oct-TyrR* and *Dop1R1* are of special interest. *Oct-TyrR* encodes an octopamine-tyramine receptor expressed in mushroom bodies [85], and *Dop1R1*, which encodes a dopamine receptor enriched in the mushroom bodies, has previously been implicated in aversive and appetitive conditioning [86], innate courtship behavior [87] and sleep-wake arousal [88]. Loss-of-function mutations of *Dop1R1* increase sleep and these effects are reversed by administration of cocaine [88]. Octopamine and tyramine act on astrocytes via the Oct-Tyr1 receptor and this activation of astrocytes can in turn modulate dopaminergic neurons [89]. Thus, we can hypothesize that combinations of octopaminergic and dopaminergic signaling in the mushroom bodies can modulate drug consumption and/or experience-dependent changes in consumption or preference following repeated exposure to cocaine or methamphetamine.

Finally, genes which were functionally validated with RNAi represent evolutionarily conserved processes. Future studies can assess whether their human counterparts play a role in variation in susceptibility to psychostimulant drug use in human populations.

## MATERIALS AND METHODS

### Drosophila stocks

The DGRP, *UAS*-RNAi and *GAL4* driver lines used are listed in Table S12. The DGRP lines are maintained in the Mackay laboratory. RNAi lines [47] were obtained from the Vienna Drosophila Resource Center and the *GAL4* driver lines from the Bloomington, Indiana Drosophila stock center. All lines were maintained on standard cornmeal/yeast/molasses medium at 25°C on a 12 hour light/dark cycle with constant humidity of 50%.

### Consumption assay

We used a two-capillary Capillary Feeder (CAFE) assay [44-46] to measure drug consumption. Briefly, five 3-5 day old flies per genotype/sex were anesthetized using CO_2_ and placed on cornmeal/yeast/molasses/agar medium one day prior to the assay. Flies were transferred without anesthesia 45 minutes prior to the assay to vials containing 4-5ml of 1.5% agar (Sigma Aldrich). Two capillaries (VWR International: 12.7 cm long, 5 µl total volume) containing 4% sucrose (Sigma Aldrich) + 1% yeast (Fisher Scientific) or 4% sucrose + 1% yeast + drug, with a mineral oil (Sigma Aldrich) overlay (to minimize evaporation), were inserted in the top of each vial. Cocaine and methamphetamine were obtained from the National Institute on Drug Abuse under Drug Enforcement Administration license RA0443159. Flies were allowed to feed for 16-18 hours with the vials placed in an enclosed plastic chamber wrapped in a plastic bag under a 12 hour light/dark cycle with constant humidity of 50%. For each experiment, an identical set of vials without flies was included in each chamber to determine evaporation loss. The capillaries were then removed and the volume of food consumed (1 mm = 0.067 µl) in each calculated as described previously [90]. The capillaries were replaced with a Drosophila activity monitor tube (TriKinetics, Inc. Waltham, MA) containing standard cornmeal/yeast/molasses medium for a recovery period of 4-6 hours. The assay was performed on three consecutive days for each vial of flies. A total of 10 replicate vials were tested for each genotype and sex.

We defined four behaviors: total amount of each solution consumed, drug preference, and change in consumption and change of preference between exposures 3 and 1. Preference was quantified in two ways: as the difference between the amount of drug and sucrose consumed (Preference A), and as this difference scaled by the total amount consumed (Preference B).

### Genetic variation in drug consumption behaviors in the DGRP

We performed four-way factorial mixed model analyses of variance (ANOVA) to partition variation in consumption in the DGRP: *Y* = *µ* + *L* + *E* + *S* + *X* + (*L* × *E*) + (*L* × *S*) + (*L* × *X*) + (*E* × *S*) + (*E* × *X*) + (*S* × *X*) + (*L* × *E* × *S*) + (*L* × *E* × *X*) + (*L* × *S* × *X*) + (*E* × *S* × *X*) + (*L* × *E* × *S* × *X*) + *ε*, where *Y* is consumption; *µ* is the overall mean; *L* is the random effect of line; *E*, *S*, and *X* are the fixed effects of exposure (day 1-3), solution (drug, sucrose), and sex (males, females); and *ε* is the residual variation between replicate vials. The main effect of L and all interaction terms with *L* are genetic factors affecting drug consumption. We also ran the same ANOVA models to compare the effects of cocaine and methamphetamine on consumption, separately for males and females. The full model for variation in change in consumption over time is *Y* = *µ* + *L* + *S* + *X* + (*L* × *S*) + (*L* × *X*) + (*S* × *X*) + (*L* × *S* × *X*) + *ε.* We assessed variation in the development of preference using the model *Y* = *µ* + *L* + *E* + *X* + (*L* × *E*) + (*L × X*) + (*E* × *X*) + (*L* × *E* × *X*) + *ε* We also assessed whether there is natural variation in the change of preference over time using the model *Y* = *µ* + *L* + *X* + (*L* × *X*) + *ε*. We also ran reduced models for each trait. All ANOVAs were performed using the PROC GLM function in SAS. We used the R function pf to assign exact *P-*values.

### Quantitative genetic analyses in the DGRP

We used the SAS PROC MIXED function to estimate variance components for each of the random effect terms in the full and reduced models. The R package lmer and lmerTest were utilized in combination with the pchisq function to assign *P*-values for the segregating genetic variation for each trait. We computed broad sense heritabilities as the sum of all genetic variance components divided by the total phenotypic variance for each model, and broad sense heritabilities of line means as the sum of all genetic variance components divided by the sum of all genetic variance components plus the environmental variance/10, where 10 is the number of replicate vials per line, sex, exposure and treatment. We computed pairwise genetic correlations as 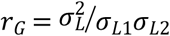, where 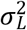 is the among line variance from the appropriate two-way factorial ANOVA and σ_*L*1_ and σ_*L*2_ are the among line standard deviations from the one-way ANOVA for each condition. We computed Pearson product-moment correlations of line means to estimate phenotypic correlations between different traits.

### Genome wide association mapping in the DGRP

We performed GWA analyses on line means for all consumption traits using the DGRP pipeline (http://dgrp2.gnets.ncsu.edu/). This pipeline accounts for effects of Wolbachia infection status, major polymorphic inversions and polygenic relatedness [43] and implements single-variant tests of association for additive effects of variants with minor allele frequencies ≥ 0.05. We tested effects of 1,891,456 DNA sequence variants on each trait.

### Network analysis

We annotated candidate genes identified by the GWA analyses using Flybase release 5.57 [56] and mapped gene-gene networks through the genetic interaction database downloaded from Flybase. We then constructed a subnetwork using Cytoscape 3.5.1 where candidate genes directly interact with each other. We evaluated the significance (α = 0.05) of the constructed subnetwork by a randomization test [91-93].

### Gene Ontology analysis

We carried out gene ontology (GO) enrichment analysis with PANTHER 11.1 (http://pantherdb.org/) [53,54].

### RNAi knockdown of gene expression

We used the binary *GAL4-UAS* system for RNAi-targeted knockdown of expression of candidate genes associated with variation in consumption of cocaine or methamphetamine with a weak ubiquitous driver (*Ubi156-GAL4*) and drivers specific for neurons (*elav-GAL4*), glia (*repo-GAL4*), mushroom bodies (*201Y-GAL4*) and dopaminergic neurons (*TH-GAL4*). We crossed 3 homozygous *GAL4* driver males to 5-7 homozygous females harboring a unique *UAS-RNAi* transgene or the progenitor control to generate F1 *GAL4-UAS-RNAi* and *GAL4* control progeny. We assessed the consumption traits exactly as described above for the DGRP lines. Differences between RNAi lines and their corresponding control lines for consumption were assessed with a fixed-effect ANOVA, separately for males and females. The full model was: *Y = µ* + *L + E* + *S* + (*L* × *E*) + (*E* × *S*) + (*L* × *S*) + (*L* × *E* × *S*) + *ε*, where *Y* denotes the mean consumption, *E* denotes the different exposures, *L* is the line (Control or RNAi), *S* denotes the different solutions (sucrose or cocaine/methamphetamine), and *ε* the error variance. Differences between RNAi lines and controls for change in consumption and preference were also assessed with fixed-effect ANOVAs. The full model for change in consumption was: *Y = µ* + *L* + *S* + (*L* × *S*) + *ε*, while the full model for preference was *Y = µ* + *L* + *E* + (*L × E*) + *ε*. All ANOVAs were run using R.

## ACKNOWLEDGMENTS

We thank Mathew J. Eddinger, Emily L. Davis, Kishan N. Patel, and Shaunaci Stevens for technical assistance. This work was supported by grants U01 DA041613 and R01 AA016560 from the National Institutes of Health to TFCM and RRHA.

## Supplementary Table Captions

**Table S1. DGRP raw consumption data.** (A) Cocaine experiment. (B) Methamphetamine experiment. F: female; M: male.

**Table S2. Analyses of variance of consumption, change in consumption, preference and change in preference of cocaine and methamphetamine.** Exposure, Sex, Solution, and their interaction are fixed effects, the rest are random. Mixed model three-way factorial ANOVAs are given for males and females, as well as reduced models by Exposure, Sex, and Solution. *E*: Exposure; *X*: Sex; *S*: Solution; *L*: DGRP Line; df: degrees of freedom; MS: Type III mean squares; F: F-ratio test; *P*: *P*-value; *σ*^2^: variance component estimate; SE: standard error; *H*^2^: Broad sense heritability. Significant *P*-values are shown in red font. (A) Cocaine experiment. (B) Methamphetamine experiment.

**Table S3. DGRP line means for all traits.** (A) Cocaine experiment. (B) Methamphetamine experiment. Means are given in mm; 1 mm = 0.067 µl.

**Table S4. Genetic and phenotypic correlations between traits.** (A) Cross-sex, cross-exposure and cross-solution genetic correlations. Significant *P*-values are indicated in red font. (B) Pair-wise phenotypic correlations. Entries in the cells are the correlation coefficients and the cell color denotes the *P*-value. Red: *P* < 0.0001; orange: *P* < 0.001; yellow: *P* < 0.01; green: *P* < 0.05; white: *P* > 0.05.

**Table S5. Results of genome wide association (GWA) analyses for consumption behaviors.** (A) Top variants (*P* < 5 ×10^-5^) and associated genes for each trait. (B) Variants and genes for the cocaine traits, the methamphetamine traits, and variants and genes overlapping between the two experiments. (C) Pathway and gene ontology enrichment analysis for the cocaine GWA analyses. (D) Pathway and gene ontology enrichment analysis for the methamphetamine GWA analyses.

**Table S6. DGRP candidate genes and human orthologs.** The references indicate which of the human orthologs have been associated with addictive phenotypes.

**Table S7. A significant genetic interaction network with no missing genes.** (A) Genes in network. (B) Pathway and gene ontology enrichment analysis.

**Table S8. Raw cocaine and sucrose consumption data for RNAi and control genotypes.** (A) *Ubi156-GAL4*. (B) *elav-GAL4*. (C) *repo-GAL4*. (D) *201Y-GAL4*. (E) *TH-GAL4*. Data are given in mm; 1 mm = 0.067 µl.

**Table S9. Raw methamphetamine and sucrose consumption data for RNAi and control genotypes.** (A) Ubi156-GAL4. (B) *elav-GAL4. (C) repo-GAL4*. (D) 201Y-GAL4. (E) *TH-GAL4*. Data are given in mm; 1 mm = 0.067 µl.

**Table S10. Analyses of variance of consumption, change in consumption, preference and change in preference of cocaine and sucrose in RNAi lines and their controls.** Fixed effect three-way factorial ANOVAs are given for males and females as well as reduced models by Exposure and Solution. E: Exposure; S: Solution; L: RNAi or control genotype; df: degrees of freedom; MS: Type III mean squares; F: F-ratio test; *P: P*-value. Significant *P*-values are shown in red font. (A) *Ubi156-GAL4*. (B) *elav-GAL4*. (C) *repo-GAL4*. (D) *201Y-GAL4*. (E) *TH-GAL4*.

**Table S11. Analyses of variance of consumption, change in consumption, preference and change in preference of methamphetamine and sucrose in RNAi lines and their controls.** Fixed effect three-way factorial ANOVAs are given for males and females as well as reduced models by Exposure and Solution. E: Exposure; S: Solution; L: RNAi or control genotype; df: degrees of freedom; MS: Type III mean squares; F: F-ratio test; *P: P*-value. Significant *P*-values are shown in red font. (A) *Ubi156-GAL4*. (B) *elav-GAL4*. (C) *repo-GAL4. (D) 201Y-GAL4*. (E) *TH-GAL4*.

**Table S12. Drosophila lines used in this study.** (A) DGRP lines. (B) RNAi lines and control genotypes. (C) *GAL4* driver lines.

**Supplementary Fig. Captions**

**Fig. S1. *P*-value summary from three-way ANOVA models of consumption for *UAS-RNAi* and control genotypes of candidate genes crossed to a weak ubiquitous *GAL4* driver (*Ubi156-GAL4*).** Red: *P* < 0.0001; orange: *P* < 0.001; yellow: P < 0.01; green: *P* < 0.05; white: *P* > 0.05.

**Fig. S2. Differences between *Ubi156-GAL4 RNAi* and control genotypes for 34 candidate genes.** (A) Cocaine preference, females. (B) Cocaine preference, males. (C) Change in cocaine preference between third and first exposures, females. (D) Change in cocaine preference between third and first exposures, males. Asterisks represent significant *L*×*S* terms (A, B) or significant *L*×*S*×*E* terms from the full ANOVA models. Exact *P*-values are given in Table S11.

**Fig. S3. Differences between *Ubi156-GAL4 RNAi* and control genotypes for 34 candidate genes.** (A) Methamphetamine preference, females. (B) Methamphetamine preference, males. (C) Change in methamphetamine preference between third and first exposures, females. (D) Change in methamphetamine preference between third and first exposures, males. Asterisks represent significant *L*×*S* terms (A, B) or significant *L*×*S*×*E* terms from the full ANOVA models. Exact *P*-values are given in Table S12.

**Fig. S4. *P*-value summary from three-way ANOVA models of consumption for *UAS-RNAi* and control genotypes of candidate genes crossed to neuronal (*elav-GAL4*) and glial (*repo-GAL4*) *GAL4* drivers.** Red: *P* < 0.0001; orange: *P* < 0.001; yellow: *P* < 0.01; green: *P* < 0.05; white: *P* > 0.05.

**Fig. S5. *P*-value summary from three-way ANOVA models of consumption for *UAS-RNAi* and control genotypes of candidate genes crossed to mushroom body (*201Y-GAL4*) and dopaminergic (*TH-GAL4*) *GAL4* drivers.** Red: *P* < 0.0001; orange: *P* < 0.001; yellow: *P* < 0.01; green: *P* < 0.05; white: *P* > 0.05.

